# Structural, functional and evolutionary analysis of wheat WRKY45 protein: A combined bioinformatics and MD simulation approach

**DOI:** 10.1101/2022.11.18.517088

**Authors:** Prashant Ranjan, Ashok Yadav, Ananta Keshari Behera, Dhiraj Kumar Singh, Premkant Singh, Ganga Prasad Singh

## Abstract

Bread wheat (Triticum aestivum L.) is the world’s second-most important cereal crop, as well as India’s. It is an allohexaploid composed of three homeologous sub-genomes (AA, BB, and DD), which is a constraint in determining the complete genome sequence. Several transcription factors have been implicated in both abiotic and biotic stress. WRKY transcription factors are among the best characterised in the context of pathogen defence mechanisms. Different members of the WRKY transcription factors have been shown to confer resistance to stress. But very little is known about the wheat WRKY transcription factors. In silico analysis of the TaWRKY45 protein was performed in the present study using several bioinformatics tools like motif scan, CD search, Netphos, NGlycos, GRAVY, and the SWISS MODEL. The study revealed that TaWRKY45 belongs to the group III family and contains hydrophilic proteins with 19 potential phosphorylation sites. TaWRKY45 protein was found to be orthologous to rice OsWRKY45 by phylogenetic analysis. The catalytic domain was analysed by motif scan which showed that TaWRKY45 has one WRKY domain and a C2-HC zinc finger motif. TaWRKY45’s structure was determined to be more stable, more constrained, more compact, and have greater potential to interact with other molecules than OsWRKY45, according to MD simulation analysis. Thus, in silico analysis of transcription factors helps study protein function, interaction, and regulatory pathways.

## Introduction

Bread wheat (Triticum aestivum L.) is one of the leading cereal crops, and India is the second largest producer of wheat in the world after China. Wheat is an allohexaploid plant consisting of three homeologous sub-genomes, AA, BB, and DD. Wheat’s large genome size (16.9 Gb) makes full genome sequencing difficult. Several Transcription Factors (TFs) are known to be involved in abiotic as well as biotic stresses [Rushton et al., 2010]. Different members of WRKY TFs have been shown to confer resistance to a variety of stresses. But very little is known about the wheat WRKY TFs [Wu et al., 2008]. WRKY TFs have a WRKY domain of ∼60 amino acids. Members of this family contain at least one conserved DNA-binding region, designated the WRKY domain, that comprises the highly conserved WRKYGQK amino acid sequence and a zinc-finger like motif. The WRKY transcription factors are categorised into three classes on the basis of the number of WRKY domains and the type of zinc-finger like motif. Group I has two WRKY domains and a C2H2 (C–X4–5–C–X22–23–H–X1–H) zinc finger. Groups II and III have one WRKY domain each. Group II has a zinc finger motif similar to Group I but Group III has a different zinc finger motif of C2HC (C–X7–C–X23–H–X–C). Group II is further divided into five subgroups based on additional amino acid motifs present outside the WRKY domain. One of these WRKY TFs involved in the positive response against Fusarium head blight and stress pathways in wheat is TaWRKY45, which confers enhanced resistance towards Fusarium graminearum [Bahrini et al., 2011]. Therefore, these particular TFs were selected for the present study to analyse their fine structure, which might help in predicting their other functions. We performed bioinformatics analysis on TaWRKY45 using various online and offline tools and also used the MD simulation technique for a comparative analysis of the structures of TaWRKY45 and OsWRKY45. We observed that the TaWRKY45 was orthologous to the OsWRKY45 by motif and phylogenetic analysis. The structural analysis of TaWRKY45 was altered as compared to OsWRKY45.

## Materials and methods

### Protein sequences

TaWRKY45’s amino acid sequence (GenBank accession ID: BAK53496) was obtained in fasta format from the NCBI database (http://www.ncbi.nlm.nih.gov/).The different WRKY sequences used for phylogenetic analysis were retrieved from NCBI.

### Analysis tools

The conserved domains of TaWRKY45 were searched using the Conserve Domain Database (http://www.ncbi.nlm.nih.gov/Structure/cdd/wrpsb.cgi) [Marchler et al., 2011]. The Protparam tool (http://web.expasy.org/protparam/) [Gasteiger et al., 2005] was used to calculate the theoretical isoelectric point (pI), molecular weight (MW), the grand average hydropathicity (GRAVY), the total number of acidic (-R) and basic (+R) residues, the alipahatic index (AI), the extinction coefficients (EC), and the instability index (II) of TaWRKY45. Netphos 2.0 server (www.cbs.dtu.dk/services/NetPhos/) [Blom et al., 1999] was used to predict the serine, threonine, and tyrosine phosphorylation sites, whereas NetNGlyc 1.0 server (http://www.cbs.dtu.dk/services/NetNGlyc/) [Lam et al 2013] was used to predict the N-Glycosylation site. Motif scan (myhits.isb-sib.ch/cgi-bin/motifscan) [Sigrist et al., 2010] was used to predict the catalytic domain site. SOPMA (http://npsa-pbil.ibcp.fr/cgi-bin/npsa_automat.pl?page=/NPSA/npsa_sopma.html) [Geourjon & Deléage., 1995] was used to predict the secondary structure of the WRKY45 protein. The automated mode of the Swiss modeller [Schwede et al., 2008] was used to predict the tertiary structure of TaWRKY45, and the structure of domain regions was only selected using the visualisation software Discovery Studio. The crystal structure of OsWRKY45 (PDB ID: 6IR8) was downloaded from the RCSB PDB. Energy minimization of structures has been done by GROMACS (Pronk et al. 2013). Further structural evaluation was done by Ramachandran. pdb sum generated a plot analysis (Laskowski et al., 2001). A phylogenetic analysis was performed in MEGA5.1 software to reveal its evolutionary significance and relationship with other WRKY TFs. [Tamura et al., 2011]. Molecular docking of DNA (Wbox-TTTGACC) and protein (TaWRKY45 and OsWRKY45) interactions was performed by a patch dock server (Schneidman et al., 2005). We used to select default parameters for docking. The crystal structure of W-box was retrieved from the RCSB PDB database, and its id was 6IR8. In Discovery Studio, visualisation of the interaction of complex systems was used. Easy structural study of DNA-protein complexes is made possible by DNApro DB, a database, structure processing pipeline, and web-based visualisation tool (Sagendorf et al., 2017). Individual nucleotide-residue interactions, DNA secondary structure, protein secondary structure, and DNA interaction moieties are all displayed on the residue contact map. The nucleotides in the DNA are represented as nodes on a graph, with the edges between them denoting backbone connections, base pairing, or base stacking. The base-pair edges show different base pairing geometries, and additional structural traits including backbone fractures, missing phosphates, and DNA strand sense are shown. Protein residues are represented as little nodes, with the shape and colour of the node signifying the secondary structure of the residue. The edges between the residue and nucleotide nodes show how the two interact with one another and which DNA molecule(s) are involved. For a particular DNA helix, the Helical Contact Map displays protein secondary structural elements (SSE) interactions along the helical axis. Concentric annuli are used to depict how SSEs interact with various DNA moieties and helicoidal coordinates—a curvilinear coordinate system determined by the axis of a helical DNA segment is used to specify the location of each depicted SSE. For a particular helix, the Helical Shape Plot depicts DNA shape parameters (such as major and minor groove widths or helical parameters) along the helix’s sequence. Furthermore, interactions between DNA and protein residues are displayed, roughly indicating how each residue at the interface interacts with the DNA sequence and the secondary structure of that residue. If only DNA shape parameters are required, protein residue interactions can be toggled off. Alternatively, they can be shown to show potential DNA shape readouts, such as the presence of positively charged residues in areas with a small minor groove width.

### MD Simulations

The equilibrium and dynamic behaviour of the TaWRKY45 and OsWRKY45 proteins were investigated using GROMACS [Poussu et al., 2004, Forloni et al., 2002]. The change in unique nature is made possible by time-dependent structural variations and corrections brought about by MD modelling once a protein mutation has been identified. GROMOS96 [Dobson., 2003] served as the force field for the GROMOS96 54a7 MD simulation. To make spc216.gro a non-exclusively equilibrated 3-point dissolvable water model in a dodecahedron, we added solvent water surrounding proteins.In this situation, the corners of the casing were kept at least 1.0 nm away from the central protein. Using the steepest descent method, steric conflicts and unstable conformations were also removed during the energy reduction procedure. NVT (constant number of particles, volume, and temperature) and NPT (constant number of particles, pressure, and temperature) ensembles are also used to further equilibrate the system. After the process achieved equilibrium, the MD run was modified to 100 ns. The data was analysed using Gromacs tools, specifically gmx rms for RMSD (Root Mean Square Deviation), RMSF (Root Mean Square Fluctuation), Rg (Radius of gyration), SASA (Solvent Accessible Surface Area), and intra H-bond. To better visualise the data, we used XM-GRACE software [Cowan & Grosdidier.,2000].

## Results

### Prediction of conserved domain and zinc-finger motif

TaWRKY45 is a protein comprised of 293 amino acid residues, and conserved domain analysis showed the presence of one WRKY domain and a C2HC type of zinc finger motif. Hence, WRKY45 is a member of the WRKY superfamily (Fig. 1), and since TaWRKY45 contained a C2HC type of zinc finger motif; it belongs to Group III (Fig. 1).

**Fig.1.**
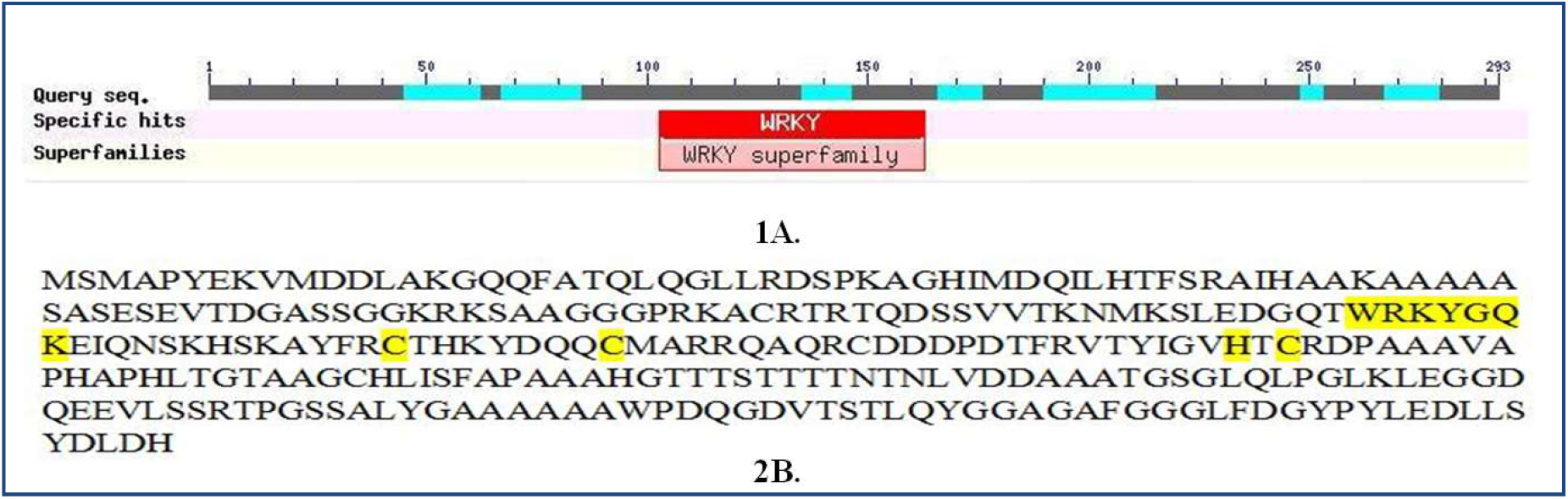
Conserve domain and Zinc finger motif analysis. 1A. NCBI conserved domain search result. 2B. Protein sequences of TaWRKY45 which has C_2_HC zinc-finger motif.

### Physico-chemical prediction of TaWRKY45

The various physico-chemical properties of the selected protein were calculated using Expasy’s protparam server. pI (Iso-electronic point), -R (negative residues) and +R (positive residues), AI, EC, II and GRAVY were determined and are provided in Table 1.

**Table 1:** Physico-chemical parameters of TaWRKY45 Protein.

| **pI** | **-R** | **+R** | **EC** | **II** | **AI** | **GRAVY** |
| --- | --- | --- | --- | --- | --- | --- |
| 6.59 | 31 | 29 | 26275 | 41.79 | 60.51 | -0.506 |

### Prediction of Hydrophobicity, Phosphorylation and N-Glycosylation sites

Hydrophobicity fluctuations were observed on a residue basis (Fig. 2). The conformation of the protein depends on its hydrophobicity(Fig. 2). The conformation of the protein depends on its hydrophobicity. There are 19 predicted phosphorylation sites in the TaWRKY45 protein, of which 9 are serine, 8 are threonine, and 2 are tyrosine (Fig. 2). Protein phosphorylation at serine, thereonine, or tyrosine residues affects a multitude of cellular signalling processes. No N-glycosylation potential site is found in TaWRKY45 (Fig. 2).

**Fig.2.**
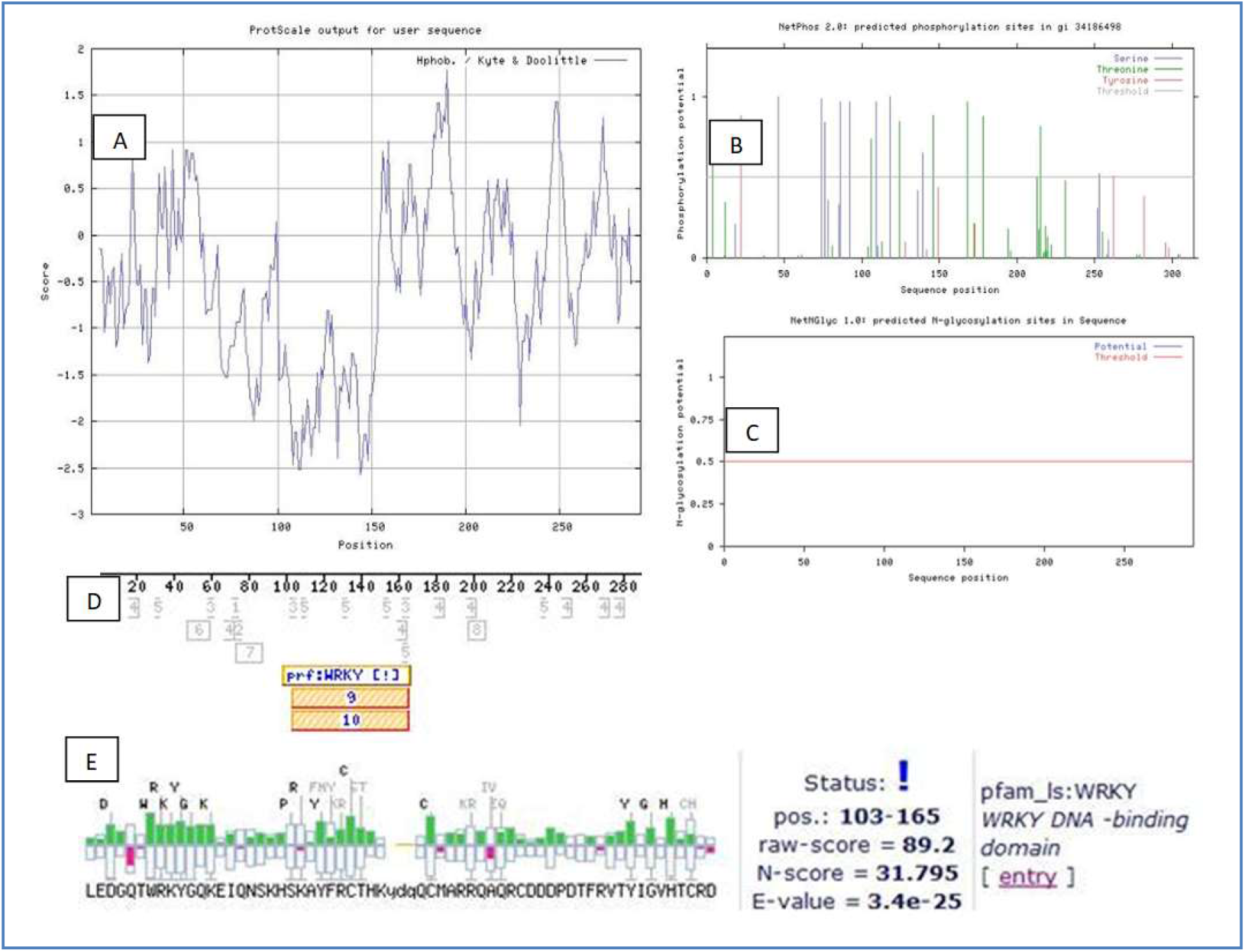
The prediction of TaWRKY45 functional aspects is followed by various protein-influencing parameters. A. Hydrophobicity plot. B & C phosphorylation and glycosylation site plots D. The position of the catalytic domain is indicated by the motif scan results. E. Graphical representation of WRKY DNA-binding domains.

### Prediction of catalytic domain arrangement

Motif scan predicted all catalytic domains present in TaWRKY45 protein (Table 2), and it also revealed the position of the catalytic domain (Fig. 2). The WRKY DNA binding region (Fig. 2), which binds specifically to the DNA sequence motif (T) (T) TGAC [CT], was also determined.

**Table 2:**
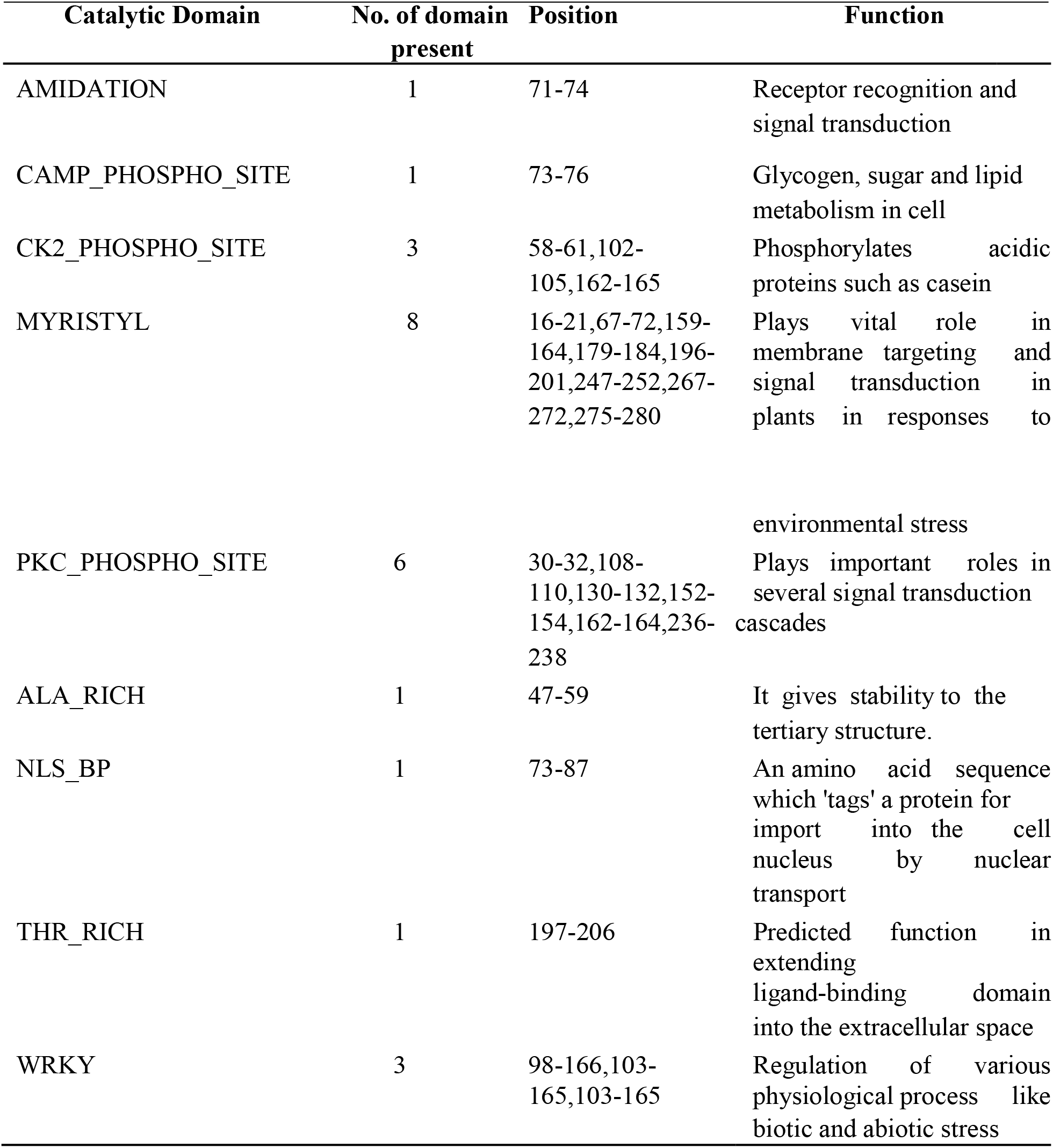
Catalytic domain arrangement in deduced amino acid sequence identified by Motif scan.

### Phylogenetic analysis of TaWRKY45

The evolutionary relationship of the TaWRKY45 sequence with other WRKY TFs (Wu et al., 2008) was visualised by constructing an unrooted phylogenetic tree using Mega 5.1 software. The TaWRKY45 protein was found to be orthologous to rice OsWRKY45 (Fig. 3).

**Fig. 3.**
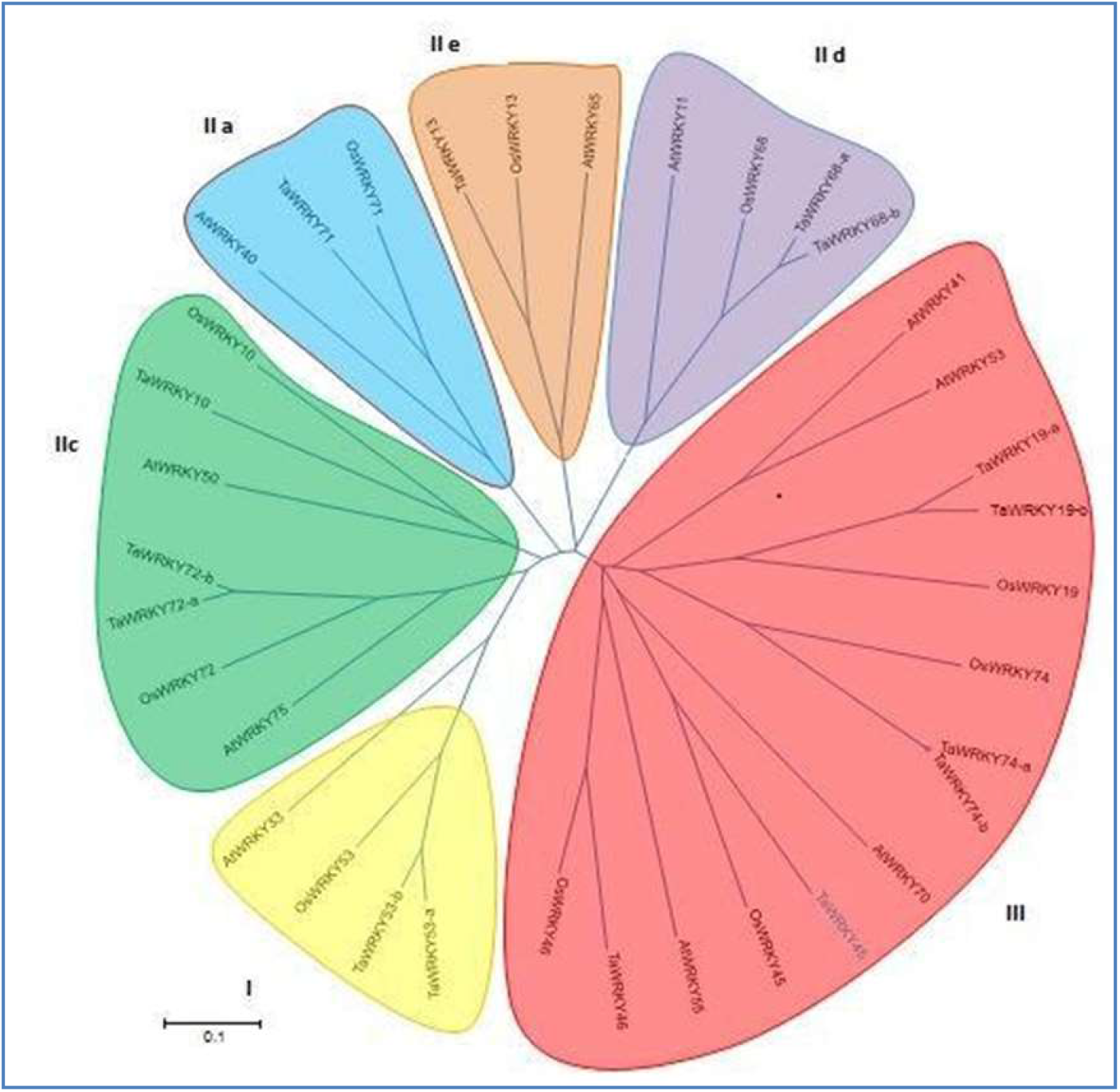
Relationships among WRKY TFs, as illustrated by the unrooted phylogenetic tree produced by MEGA5.1. The scale represents the branch length. TaWRKY45 protein sequence clusters with other WRKY TFs in group III.

### Secondary and Tertiary structure prediction of WRKY45

Secondary structure prediction was performed by SOPMA for calculating protein structures such as alpha helices and beta strands (Table 3). The Swiss model predicted the TaWRKY45 tertiary structure and retrieved the orthologous OsWRKY45 structure from the RCSB PDB, which consists of four antiparallel sheets and three random coils (Fig. 4).

**Table 3:** SOPMA results of the TaWRKY45 protein.

| **Parameters** | **Total contents of parameters** | **% contents of parameters** |
| --- | --- | --- |
| Alpha helix | 98 | 33.45% |
| Extended strand | 36 | 12.29 |
| Beta turn | 14 | 4.78% |
| Random coil | 145 | 49.49% |

**Fig 4.**
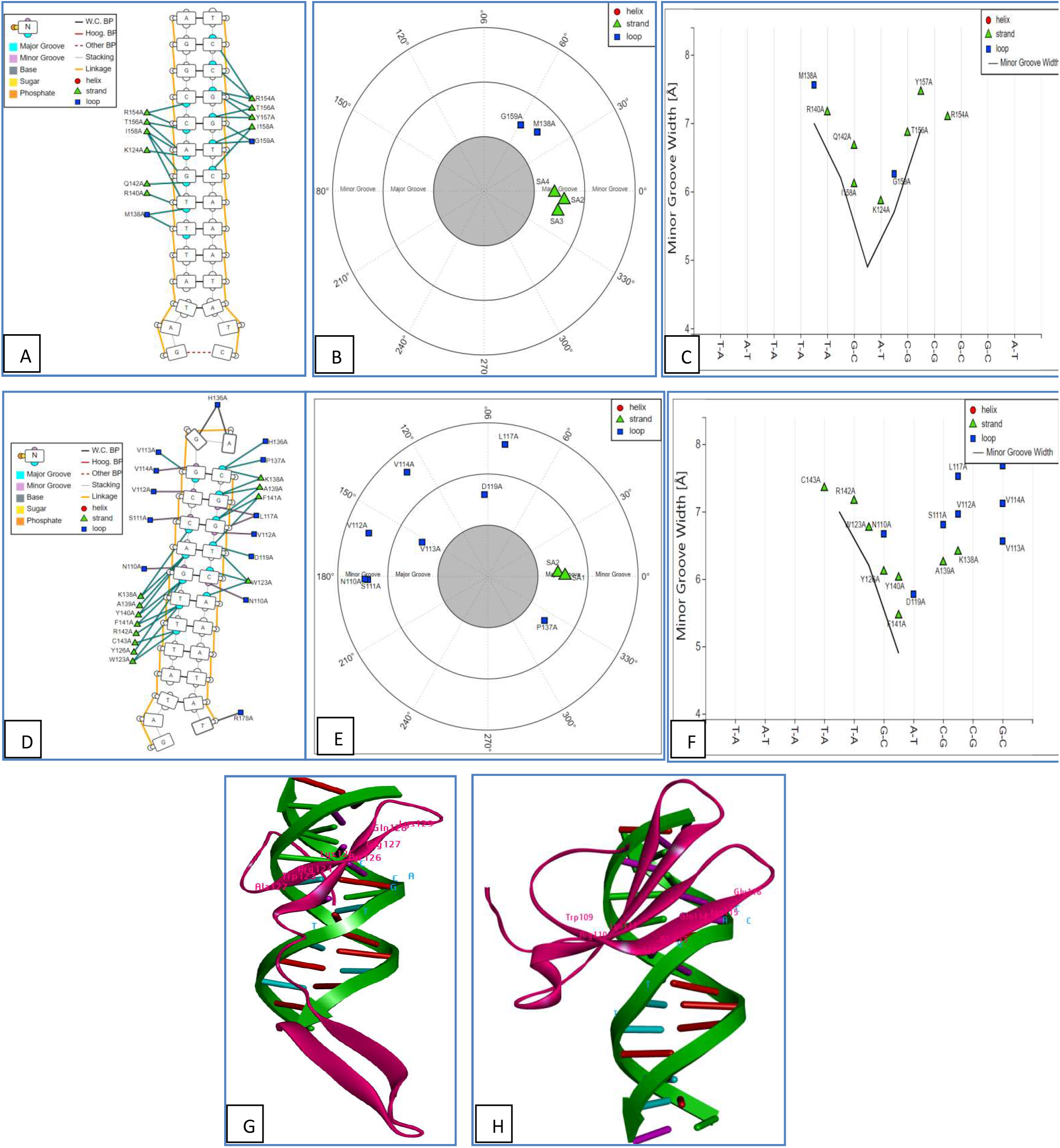
TaWRKY45 and OsWRKY45 DNA proteins interface with the W-box. A and D displayed a residue contact map, B and E a helical contact map, C and F a helical shape overlay, and G and H indicated that OsWRKY45 and TaWRKY45 were W-box complexes respectively. A, B and C indicated Protein DNA interfaces of TaWRKY45 and D, E and F showed Protein DNA interfaces of OsWRKY45. DNA Protein interaction noted higher in OsWRKY45 with W-box as compared to TaWRKY45. TaWRKY45 interface analysis showed number of residues on strands and loops participated in interactions were less as compared to OsWRKY45 interface analysis. TaWRKY45 residues (B) showed interaction mainly in the major groove area, while in OsWRKY45; more residues (E) involved both the major and minor grooves. The contacts between DNA and protein residues are shown (C&F), approximately illustrating the locations of each interface residue’s interactions with the DNA sequence and the residue’s secondary structure. G&H depicted the interaction complex in 3D.

### Protein DNA interaction prediction

We looked at studies of interactions between WRKY45 (TaWRKY45 & OsWRKY45) and Wbox and came to the conclusion that TaWRKY45 had a lesser binding score with Wbox than OsWRKY45. With Wbox, a binding score of 9888 was recorded against TaWRKY45. However, the Wbox for OsWRKY45 was 14336. The orientation of interaction was depicted in Fig.4.

### MD Simulation

After 30 ns of simulation, the RMSD plot graph got equilibrated, and the average RMSD value of TaWRKY45 was shown to be 1.8 nm, while ∼2.75 nm was noted in OsWRKY45. RMSF value fluctuation was observed in TaWRKY45, i.e., 0.2-0.75 nm, while in OsWRKY45, 0.25 to 1.9 nm. The Rg value in TaWRKY45 equilibrates after 10 ns, whereas it equilibrates after 50 ns in OsWRKY45. After equilibration was achieved, the average Rg value for TaWRKY45 was ∼1.75 nm, while more Rg was observed in OsWRKY45, i.e., ∼2 nm. After 40 ns, the SASA plot gets equilibrated in both Ta and Os WRKY45, and the average SASA was observed at ∼90 nm2 in TaWRKY45 while ∼105 nm2 was observed in OsWRKY45. The fluctuation of h-bonds was observed to be higher in TaWRKY45 as compared to OsWRKY45. The MD simulation plots were depicted in Figs. 5 & 6.

**Fig. 5.**
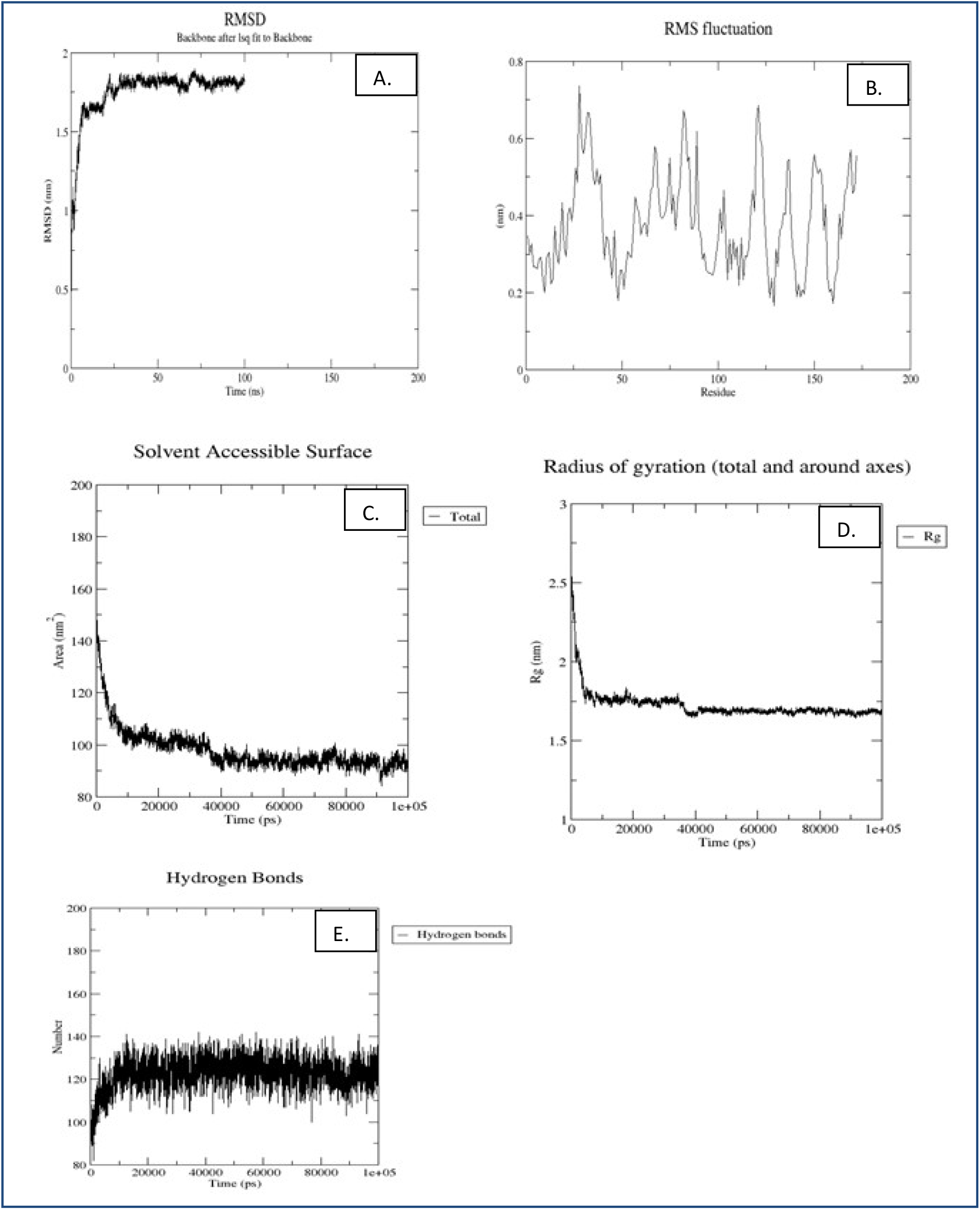
Molecular simulations results of TaWRKY 45 at 100 ns time periods. (A) RMSD plots. (B) RMSF. (C) Solvent accessible surface area (SASA). (D) Radius of gyration (Rg). (E) Intramolecular hydrogen bonding.

**Fig. 6.**
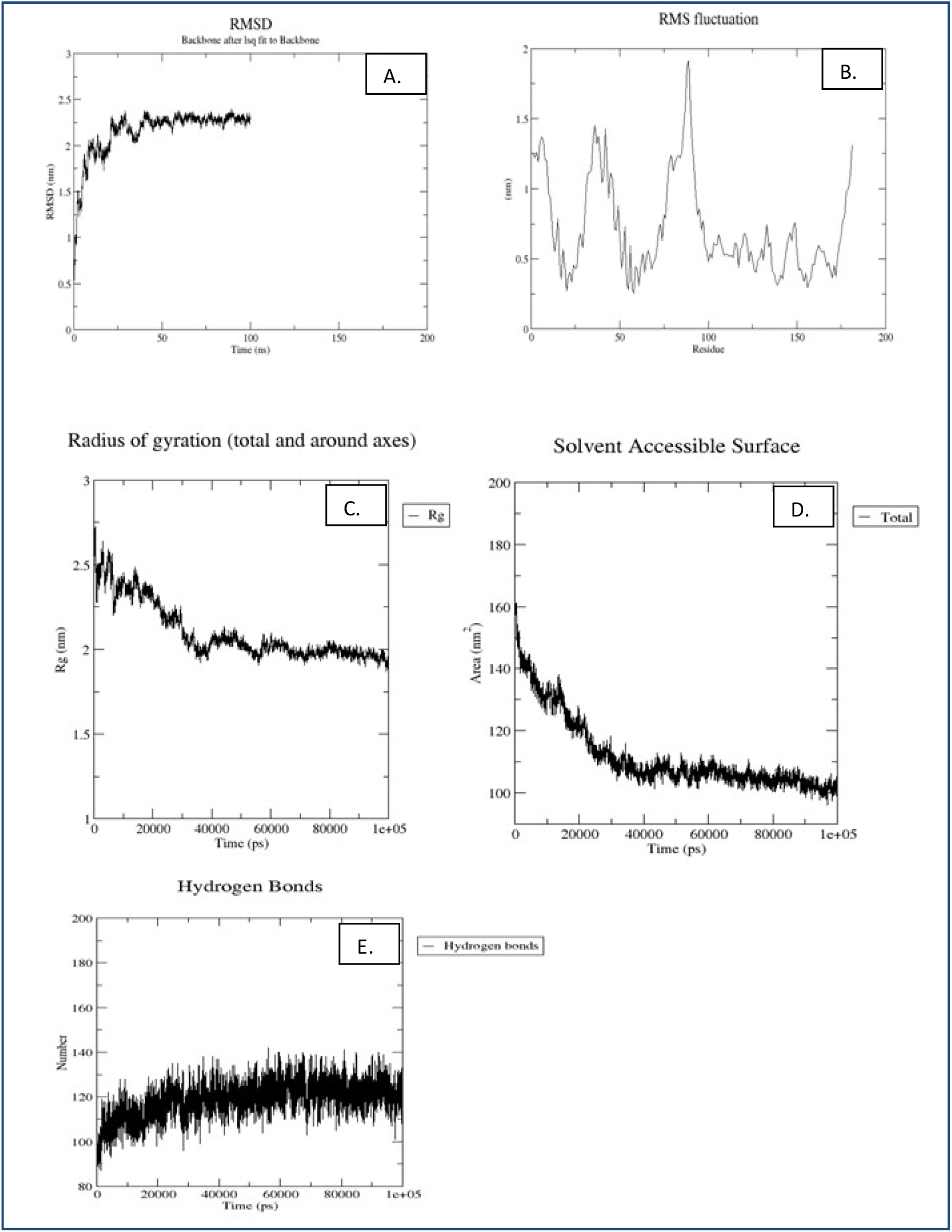
Molecular simulations results of TaWRKY 45 at 100 ns time periods. (A) RMSD plots. (B) RMSF plot. (C) Radius of gyration (Rg). (D) Solvent accessible surface area (SASA). (E) Intramolecular hydrogen bonding.

## Discussions

According to research, plants use the WRKY proteins for a variety of purposes. The WRKY proteins, according to Chen and Chen (2002), control how plants react to physical injury and pathogen assault. Furthermore, the WRKY proteins are implicated in a number of protective signal transduction pathways that develop disease resistance and response (Xu et al., 2006). TaWRKY45 transcription factor appears to contribute positively to wheat’s resistance to FHB (Fusarium Head Blight) and is engaged in the defensive response to a Fusarium assault. The expression of genes linked to numerous disease resistance and stress response pathways in wheat may also be regulated by TaWRKY45 (Bahrini et al 2011). The stress-induced OsWRKY45 gene is overexpressed, which improves disease resistance and drought tolerance (Qiu & Yu et al 2008). WRKY proteins are classified into three groups based on their structures: group I, group II and group III (Eulgem et al. 2000). WRKY45 falls into group III, previously considered to be a group associated with the defence response in Arabidopsis (Kalde et al. 2003). Here, we have performed an in silico analysis of TaWRKY45 to understand its structural, functional, and evolutionary significance.

TaWRKY45 is orthologous to OsWRKY45 and belongs to group III, according to motif and phylogenetic analyses. The physicochemical parameter analysis highlights the functional aspects of the protein. At a pI of 6.59, TaWRKY45 protein becomes the least soluble in nature.TaWRKY45 has a 31 and a 29 for its acidic and basic residues, respectively. The protein’s property turns acidic as a result of the abundance of acidic residues. The extinction coefficient provides information on how much light at a specific wavelength is absorbed by a protein. TaWRKY45 is unstable in a test tube, according to the instability index value, which is greater than 40. The protein’s 60.51 aliphatic index demonstrated the thermostability of globular proteins. TaWRKY45 has a GRAVY value of -0.506, which denotes that it is a hydrophilic protein. TaWRKY45 showed a lower binding score with Wbox than OsWRKY45, according to investigations of interactions between WRKY45 (TaWRKY45 & OsWRKY45) and Wbox. A binding score of 9888 was obtained using Wbox when compared to TaWRKY45. The Wbox for OsWRKY45 was 14336. When compared to OsWRKY45, TaWRKY45 may be less active in defence and have a higher stress response. TaWRKY45 has many catalytic domains, according to motif analysis, including AMIDATION, CAMP PHOSPHO SITE, CK2 PHOSPHO SITE, MYRISTYL, PKC PHOSPHO SITE, ALA RICH, NLS BP, THR RICH, and WRKY, which aid in the regulation of the protein’s multiple functions.

The MD simulation is conducted for 100 ns to investigate the dynamic behaviour and to compare the structural stability of the TaWRKY45 protein structure to that of the OsWRKY45 protein structure. To study the functional and structural influence of TaWRKY45 on the orthologous OsWRKY45 protein, a variety of parameters were studied throughout the simulation trajectory, including RMSD, Rg, RMSF, SASA, and the total number of intra-molecular hydrogen bonds in the protein. In comparison to OsWRKY45, TaWRKY45’s C-alpha chain atoms’ RMSD and RMSF revealed substantial changes in both flexibility and stability. The RMSD value of TaWRKY45 (1.8 nm) was less than that of OsWRKY45 (∼2.75 nm). TaWRKY45 was found to be more stable than OsWRKY45. However, the RMSF fluctuation value was higher in OsWRKY45 (0.25 to 1.9 nm) than TaWRKY45 (0.2-0.75 nm), suggesting that the structure of TaWRKY45 may be more restricted than OsWRKY45. According to the RG and SASA analyses, Ta and Os WRKY45 showed significant variation. The fact that the Rg value was observed lower in TaWRKY45 (∼1.75 nm) than OsWRKY45 (∼2 nm) indicates TaWRKY45 was more compact than OsWRKY45. The SASA value was higher in OsWRKY45 (∼105 nm2) than TaWRKY45 (∼90 nm2). It implies that the structure of TaWRKY45 is less accessible than OsWRKY45, which might impact their ability to interact with other molecules. Changes in the overall number of intramolecular H-bonds may have an impact on the stiffness of proteins and the interactions between them.

### Conclusion

TaWRKY45 consists of 293 amino acid and belongs to Group III WRKY transcription factor family. It contains one WRKY domain and a C_2_HC-type zinc finger motif. *In silico* analysis showed that TaWRKY45 is a hydrophilic protein with 19 phosphorylation sites. TaWRKY45 contains several catalytic domains, including a DNA-binding domain. Phylogenetic analysis showed TaWRKY45 to be orthologous to OsWRKY45. MD simulation analysis reveals that the structure of TaWRKY45 was found to be more stable, more restricted, more compact, and have more potential to interact with other molecules compared to OsWRKY45. *In silico* analysis of TaWRKY45 helps us study its function, interaction with the DNA binding domain, and role in different regulatory pathways.

